# Multistep changes in amyloid structure that are induced by cross-seeding on a rugged energy landscape

**DOI:** 10.1101/2020.07.19.211284

**Authors:** Keisuke Yuzu, Naoki Yamamoto, Masahiro Noji, Masatomo So, Yuji Goto, Tetsushi Iwasaki, Motonari Tsubaki, Eri Chatani

**Affiliations:** Graduate School of Science, Kobe University, 1-1 Rokkodai, Nada, Kobe, Hyogo 657-8501, Japan; School of Medicine, Jichi Medical University, 3311-1 Yakushiji, Shimotsuke, Tochigi 329-0498, Japan; Institute for Protein Research, Osaka University, 3-2 Yamadaoka, Suita, Osaka 565-0871, Japan; Global Center for Medical Engineering and Informatics, Osaka University, 2-1 Yamadaoka, Suita, Osaka 565-0871, Japan; Biosignal Research Center, Kobe University, Rokkodai, Nada, Kobe, Hyogo 657-8501, Japan

**Keywords:** amyloid, insulin, protein misfolding, protein aggregation, protein structure, cross-seeding, iodine staining, metastable state

## Abstract

Amyloid fibrils are aberrant protein aggregates associated with various amyloidoses and neurodegenerative diseases. It is recently indicated that structural diversity of amyloid fibrils often results in different pathological phenotypes including cytotoxicity and infectivity. The diverse structures are predicted to propagate by seed-dependent growth, which is one of the characteristic properties of amyloid fibrils. However, much remains unknown regarding how exactly the amyloid structures are inherited to subsequent generations by seeding reaction. Here, we investigated the behaviors of self- and cross-seeding of amyloid fibrils of human and bovine insulin in terms of thioflavin T fluorescence, morphology, secondary structure, and iodine staining. Insulin amyloid fibrils exhibited different structures depending on species, and each of which replicated in self-seeding. In contrast, gradual structural changes were observed in cross-seeding, and a new-type amyloid structure with distinct morphology and cytotoxicity was formed when human insulin was seeded with bovine insulin fibrils. Remarkably, iodine staining tracked changes in amyloid structure sensitively, and singular value decomposition (SVD) analysis of the UV-Vis absorption spectra of the fibril-bound iodine has revealed the presence of one or more intermediate metastable states during the structural changes. From these findings, we propose a propagation scheme with multistep structural changes in cross-seeding between two heterologous proteins, which is accounted for as a consequence of the rugged energy landscape of amyloid formation.

## Introduction

Protein misfolding induces aberrant aggregation, and often results in the formation of amyloid fibrils that are associated with more than 40 amyloidoses and neurodegenerative diseases, including Alzheimer’s and Parkinson’s diseases (1–4). Although needle-like morphology and cross-β structure are common basic properties of amyloid fibrils, it has been revealed that amyloid fibrils show microscopic structural diversity (4–6). Differences in amyloid structure are observed even when amino acid sequence is identical or very similar, and details of these different structures have recently been clarified at an atomic level owing to the progress of cryoelectron microscopy (cryo-EM) (7,8) and solid-state NMR spectroscopy (9,10). Variations of amyloid structure often lead to various physicochemical properties such as morphology (11,12) and structural stability (13). Moreover, it has also been suggested that such structure-dependent properties of amyloid fibrils exert diverse pathological effects in vivo (7,8,14–16) and researches on relationships between amyloid fibril structure and its phenotype have been conducted vigorously. In prion disease, which has been revealed as an infectious amyloidosis, it is proposed that the diverse structures of prion fibrils result in different transmissibility (13). Amyloid conformations are thus an important factor for determining various physiological phenotypes, and now attract much attention for understanding pathology based on protein structure.

According to a nucleation-dependent polymerization, which is the basic model of amyloid assembly, the growth of amyloid fibrils is accelerated by the presence of fibril seeds (17). With this mechanism, diverse structures of amyloid fibrils are proposed to proliferate in vivo and therefore to play an important role in the transmission and progression of diseases. Typically, seed-dependent elongation with the same proteins, i.e., self-seeding, occurs most preferably, and in this case, the structure of seeds is replicated robustly during elongation (18). As a detailed molecular picture, the dock-lock model has been provided (19). On the other hand, cross-seeding is less likely to proceed than self-seeding, although it occurs when proteins can cross a species barrier, as seen between variants of the same protein (20–22), and additionally between heterologous proteins in some cases (23,24). Several studies about the propagation of amyloid structures in cross-seeding have been conducted so far, and in an early report on prion protein variants by Jones and Surewicz, it was demonstrated that the morphology and secondary structure of seed fibrils were inherited exactly when they were used as a template in the cross-seeding of proteins from other species (20). In contrast, recent studies have revealed that cross-seeding sometimes produced amyloid fibrils different from those used as seeds with respect to secondary structure and cytotoxicity (25–27). The latter observations suggest that the complete replication of seed fibrils is not always achieved in cross-seeding, although detailed mechanisms remain unclear.

In this study, we investigated self- and cross-seeding of human and bovine insulin. Insulin is a hormone protein associated with glucose metabolism and consists of two polypeptide chains, A-chain (21 residues) and B-chain (30 residues), which are cross-linked by two interchain disulfide bonds. It is one of the best models for studying amyloid formation mechanism because it forms amyloid fibrils readily in vitro (28). Between human and bovine insulin, amino acid sequences differ in three residues (two in A-chain and one in B-chain) (Figure 1A), while their native structures are almost the same (21). Since these amino acid sequences are close enough to cross species barriers with each other, they are good models for investigating interspecies cross-seeding. We tracked preservation or change of amyloid fibril structure in the process of repetitive self-or cross-seeding of human and bovine insulin by using thioflavin T (ThT) fluorescence, atomic force microscope (AFM), Fourier transform infrared (FTIR) spectroscopy and iodine staining. In addition, the cytotoxicity of amyloid fibrils was also analyzed using cell lines as one of the indicators of phenotype.

**Figure 1.**
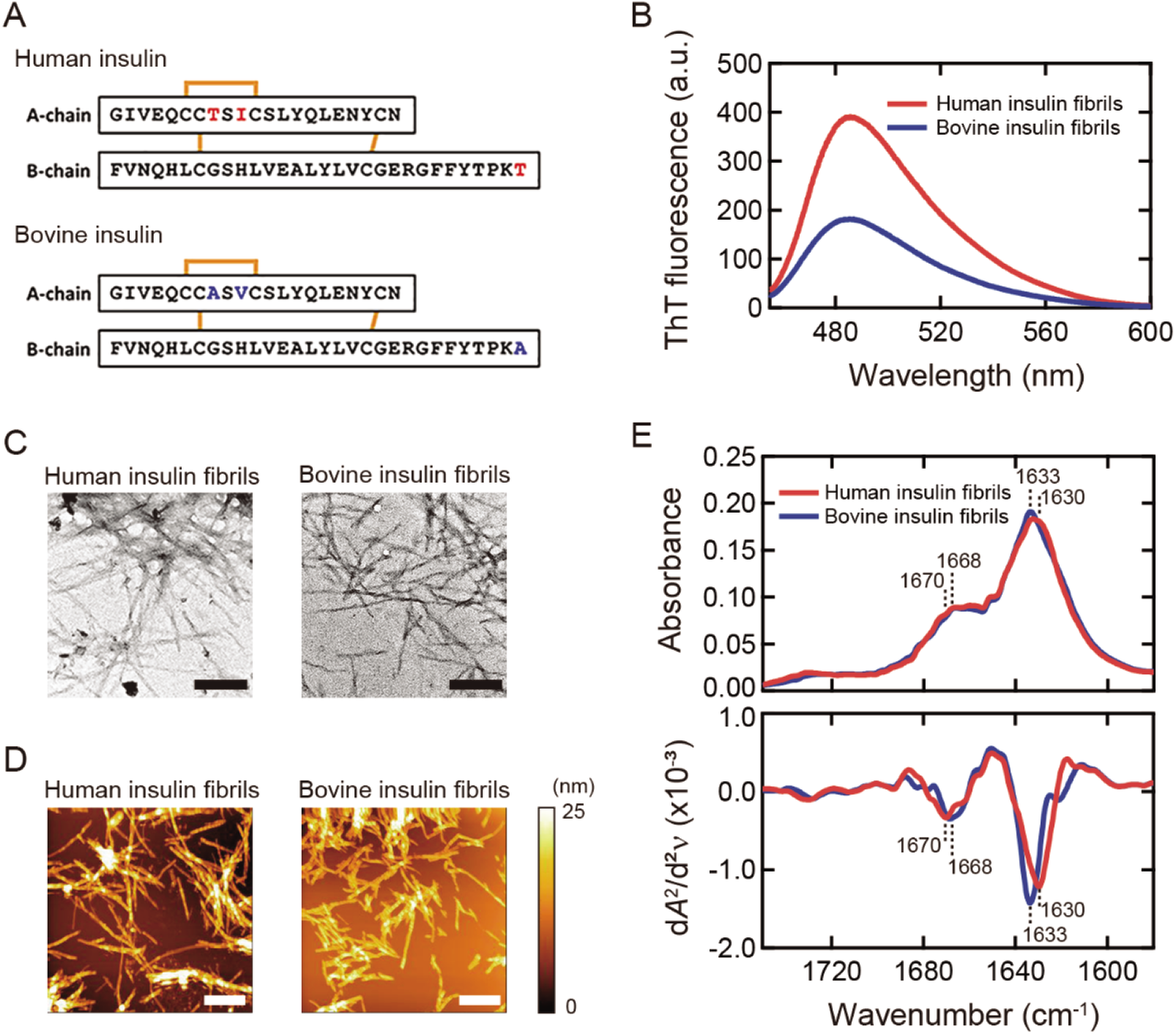
Basic properties of human and bovine insulin amyloid fibrils. (A) Amino acid sequences of human and bovine insulin. The red and blue characters represent different residues between human and bovine insulin, respectively. (B) ThT fluorescence spectra. (C) TEM images. The black scale bars represent 200 nm. (D) AFM images. The white scale bars represent 1 μm. The color scale on the right represents the height of the sample. (E) ATR-FTIR absorption spectra (upper panel) and their second derivatives (lower panel) in amide I region. The positions of two main peaks of β-sheet and β-turn structures, which were determined from the analysis of local minima of the second derivatives, are indicated on each spectrum.

The results of the measurements with several different techniques have demonstrated that human and bovine insulin amyloid fibrils adopted different structures even though only three residues are different. While both of these amyloid structures were replicated by self-seeding, they changed gradually when cross-seeding was performed. Notably, a new-type fibril structure with distinct morphology and cytotoxicity was observed when human insulin was cross-seeded with bovine insulin seeds. Here, iodine staining, a coloring reaction through the formation of polyiodide ions on the surface of amyloid fibrils (29), worked as a sensitive probe for reporting changes in amyloid structures. The UV-Vis absorption spectra of iodine-stained amyloid fibrils showed complicated changes, and by introducing singular value decomposition (SVD) analysis, intermediate metastable states have been identified in the process of cross-seeding.

## Experimental procedures

### Materials

Recombinant human insulin was purchased form Wako Pure Chemical Industries, Ltd. (Osaka, Japan). Bovine insulin was purchased from Sigma-Aldrich Co. Ltd. (St. louis, MO). Iodine, potassium iodine, and ThT were purchased from Wako Pure Chemical Industries, Ltd. HCl solution and NaCl were purchased from Nacalai Tesque, Inc. (Kyoto, Japan).

### Formation of insulin amyloid fibrils

Human and bovine insulin were dissolved in 25 mM HCl and whose concentrations were determined using an absorption coefficient of 1.08 cm^-1^ (mg/mL)^-1^ at 276 nm (30). Spontaneous fibrillation of human and bovine insulin was performed by heating; a sample solution containing 2.0 mg/mL insulin, 25 mM HCl and 100 mM NaCl was incubated at 65 °C for 24 h without agitation. For seeding experiments, seeds were obtained by sonicating pre-formed fibrils with 20 pulses with 1 sec pulse length at a power level of 2.0 W. Seeding reaction was essentially performed under the same solvent conditions as used for the spontaneous fibrillation except for the addition of 5% (v/v) seeds. The reaction temperature was set at 37 °C to prevent spontaneous fibrillation of native insulin and the incubation time was set for 18 h for each reaction.

### ThT fluorescence assay

To assay the formation of amyloid fibrils, ThT fluorescence assay was performed with fundamentally the same protocol as previously reported (31). In the assay, 3 μL of a sample solution was mixed with 1.5 mL of 5 μM ThT in 50 mM Gly-NaOH buffer (pH 8.5). After 1 min incubation, ThT fluorescence intensity at 485 nm was measured using an excitation wavelength of 445 nm with a RF-5300 PC spectrofluorometer (Shimadzu Corporation, Kyoto, Japan).

### Attenuated total reflectance (ATR)-FTIR spectroscopy

ATR-FTIR spectra were recorded with a Nicolet iS5 FT-IR equipped with an iD5 ATR accessory (Thermo Fisher Scientific, Tokyo, Japan). Each fibril sample was sonicated before measurements with the same protocol as used for the preparation of seeds to improve dispersibility. 3.5 μL of a sample solution was spotted onto the surface of a diamond crystal and then dried using compressed air. FTIR spectra were measured by collecting 128 interferograms with a resolution of 4 cm^-1^. A spectrum of atmospheric air was used as a reference spectrum to subtract the contributions of water vapor and carbon dioxide.

### Iodine staining of amyloid fibrils

Iodine staining of amyloid fibrils was performed as previously reported (29). Iodine solution containing 24 mM KI and 3.2 mM I2 was initially prepared, and each sample of 2.0 mg/mL insulin fibrils was sonicated with the same protocol as used for the preparation of seeds to improve dispersibility. Then 0.50 mg/mL insulin fibrils containing 0.9 mM KI and 0.12 mM I2 in 25 mM HCl were prepared by mixing the fibril sample and the iodine solution. After 1 min incubation, the color of the iodine-stained insulin fibrils was analyzed by UV-Vis spectroscopy. UV-Vis absorption spectra were recorded with a Jasco V-650 spectrometer (JASCO, Tokyo, Japan) using a quartz cell with a 1 cm optical length.

### Atomic force microscopy (AFM)

10 μL of a sample solution was loaded to a freshly cleaved mica plate, left for 1 min, and then rinsed using 200 μL of distilled water. Residual solution was removed with filter paper and left to dry overnight. AFM images were obtained using a dynamic force mode with Probestation NanoNavi II/IIe (Hitachi High-Tech Science, Tokyo, Japan). The scanning tip used was an OMCL-AC160TS-C3 micro cantilever (Olympus Corporation, Tokyo, Japan; spring constant = 21–37 N/m, resonance frequency = 270–340 kHz), and the scan rate was set to 0.5 Hz with the recording of 256 × 256 points per image.

### Transmission electron microscopy (TEM)

Samples were diluted 10-fold with distilled water and 5 μL of a sample solution was spotted onto a collodion-coated grid (Nisshin EM Co., Ltd., Tokyo, Japan). After 1 min, the solution on the grid was removed with filter paper. Then 5 μL of 5% (w/v) hafnium chloride was spotted onto the grid, left for 1 min, the solution was removed in the same manner. TEM images were obtained using an H-7650 transmission microscope (Hitachi High-Technologies Corporation, Tokyo, Japan) with a voltage of 80 kV.

### Cytotoxic assay

Cell viability was determined using a 3-(4,5-dimethylthiazol-2-yl)-2,5-diphenyltetrazolium bromide (MTT) cell count kit (Nacalai Tesque, Kyoto, Japan), which is based on the conversion of tetrazolium salt by mitochondrial dehydrogenase to a formazan product. PC12 cells were maintained in Dulbecco’s Modified Eagle’s medium (DMEM) containing 10% fetal bovine serum, 5% horse serum, penicillin (100 U/mL), and streptomycin (100 μg/mL) in 5% CO_2_ at 37 °C. The cells were plated in 96-well plates at a density of 20,000 cells/well and grown overnight. The cells were incubated in 110 μL medium in the absence and presence of insulin samples diluted with phosphate buffered saline (PBS) to indicated concentrations. After 24 h incubation, 10 μL of MTT reagents were added and incubated for 4 h in 5% CO_2_ at 37 °C. The reaction was stopped by adding 100 μL of solubilization solution (40 mM HCl in 2-propanol). The plates were read with a Multiskan FC microplate reader (Thermo Fisher Scientific Inc., Waltham, MA) at 570 nm. Each sample was assayed in triplicate and the cell viability was calculated using the value obtained by the addition of PBS alone as a reference.

## Results

### Formation of human/bovine insulin amyloid fibrils

Human and bovine insulin amyloid fibrils were first prepared by heating under acidic conditions, as used in earlier studies (32–34). After the completion of the reaction, more than 99 % of insulin protein was converted to fibrils for both species, which was confirmed by measuring the concentration of residual insulin monomer in the supernatant of the samples after centrifugation. When amyloid fibrils were mixed with ThT, human insulin fibrils showed a higher and slightly red-shifted fluorescence spectrum compared to bovine insulin fibrils (Figure 1B). In light of the previous suggestion that the ThT fluorescence properties vary depending on the binding mode of ThT on amyloid fibrils (35), the difference in ThT fluorescence spectrum implies that the structures of amyloid fibrils are different between these two species.

We further analyzed the morphology and secondary structure of these amyloid fibrils. TEM images exhibited needle-like morphology in both fibrils and there was no significant difference in their appearance (Figure 1C). On the other hand, AFM images revealed a difference in thickness between human (12.5 ± 2.2 nm; *n* = 20) and bovine (7.2 ± 1.4 nm; *n* = 20) insulin fibrils (Figure 1D). Furthermore, ATR-FTIR spectra in amide I region showed a subtle but significant difference in shape between the two types of insulin fibrils (Figure 1E; upper panel). The second derivative spectra showed the difference more clearly in the wavenumber of minima at around 1630 cm^-1^ corresponding to β-sheet structure (36), in agreement with the previous report by Surmacz-Chwedoruk et al (21). In addition, differences in the position of minimum at around 1670 cm^-1^ corresponding to β-turn (37) were observed (Figure 1E; lower panel). Far-UV CD spectra also showed slightly different spectral shapes between human and bovine insulin fibrils while those in native structure were quite similar (Figure S1). It was suggested that, while both of the human and bovine insulin amyloid fibrils are constructed from cross-β structure in common, there are subtle conformational differences in β-sheet and β-turn.

### Color formation of human/bovine insulin amyloid fibrils by iodine staining

We next investigated the color formation of insulin fibrils by iodine staining, whose binding modes have been proposed to vary depending on surface topology of β-pleated sheets and their edges (29). When mixed with an iodine solution, human and bovine insulin fibrils exhibited different colors (Figure 2A), giving different shapes in the UV-Vis absorption spectra mainly in a visible region (400–750 nm) (Figure 2B, C). Iodine-stained human insulin fibrils had a major absorption at around 560 nm (Figure 2B), whereas iodine-stained bovine insulin fibrils had one at around 480 nm (Figure 2C). This observation indicates that iodine staining can detect differences in fibril structure sensitively.

**Figure 2.**
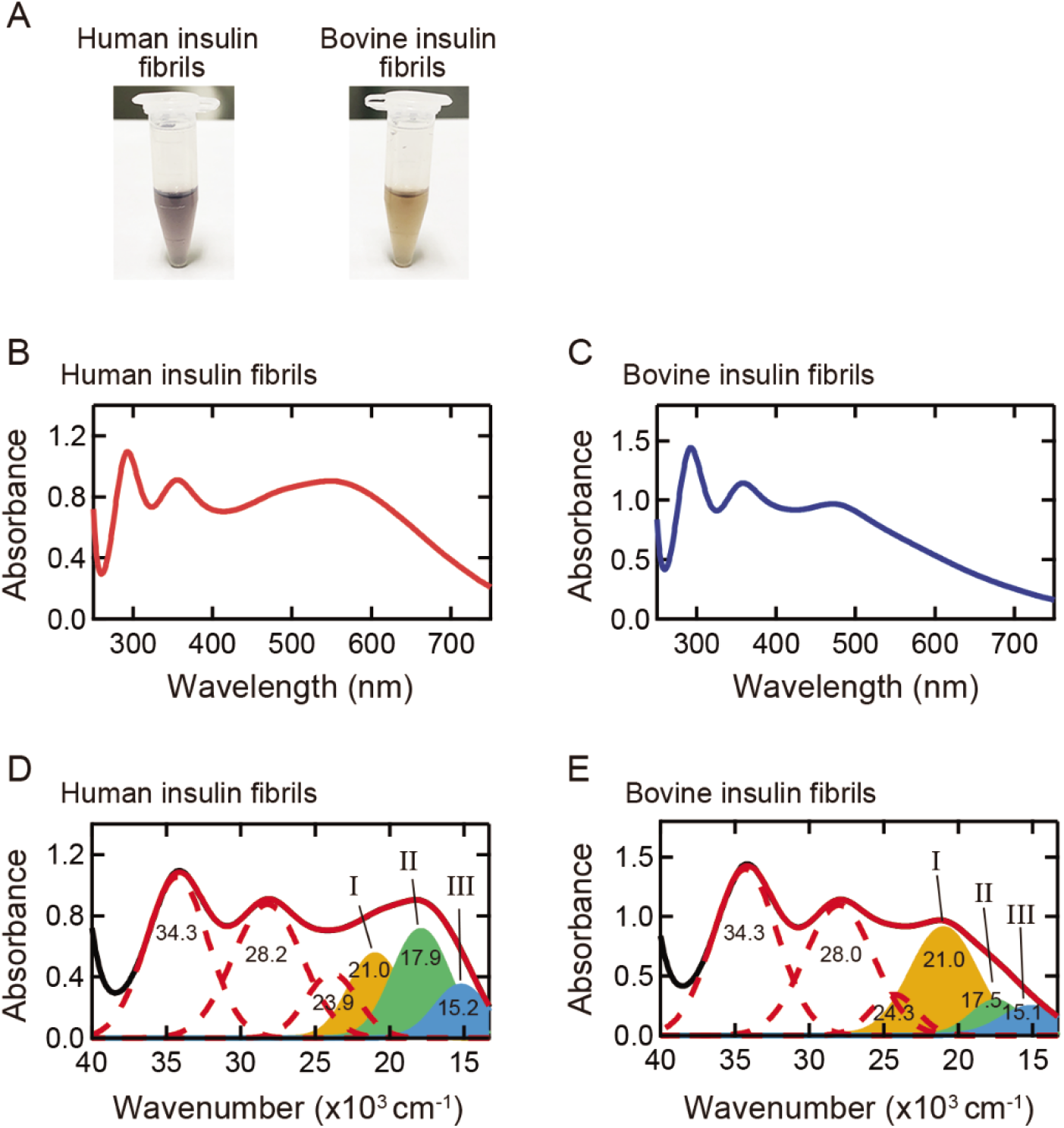
Color properties of iodine-stained insulin amyloid fibrils. (A) Photographs of iodine-stained human and bovine insulin fibrils. (B, C) UV-Vis absorption spectra of iodine-stained human (B) and bovine insulin fibrils (C). For each absorption spectrum, the spectrum of unstained fibrils was subtracted to obtain net spectrum derived from iodine molecules. (D, E) Deconvolution of UV-Vis absorption spectra shown in (B) and (C). Note that the spectra were replotted against wavenumber and the deconvolution was performed by using eq. 1. Black and red solid lines represent the experimental and reproduced spectra, respectively, and the experimental spectra are the same as shown in (B) and (C). Six Gaussian peaks are represented with red dashed lines and filled areas, which correspond to absorption components obtained by the deconvolution.

To evaluate the spectral difference in more detail, we performed spectral deconvolution and compared absorption bands constructing the experimental spectra. Each spectrum was replotted against wavenumber, and spectral deconvolution was performed using a sum of Gaussian distributions according to the following model function;

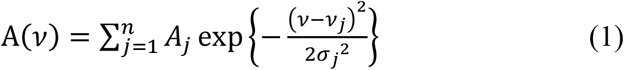

where *A_j_* and *ν_j_* represent amplitude and center wavenumber of the *j*th absorption band, respectively, and *σ_j_* is a coefficient related to the full width at half-maximum (FWHM) of the *j*th absorption band, as is described as 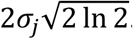. For stable and accurate convergence, two center wavenumbers and one FWHM of the absorption band parameters were fixed with reference to the local minimum and its distance to the inflection point in the second derivative spectrum.

As a result, the absorption spectra were reproduced with six Gaussians and the fitted curves reproduced the experimental spectra well (Figure 2D, E). Our previous study revealed that three absorption bands in a higher wavenumber at around 34 × 10^3^, 28 × 10^3^ and 24 × 10^3^ cm^-1^ are assigned to I_3_^-^ and I_2_ molecules (dashed lines in Figure 2D, E), and the remaining three absorption bands in a lower wavenumber at around 21 × 10^3^, 18 × 10^3^ and 15 × 10^3^ cm^-1^ are assigned to polyiodide ions larger than I3’ (filled bands in Figure 2D, E) (29). Since the I_3_^-^ and I_2_ absorption bands are observed in the iodine solution itself, the color formation by the iodine staining could be explained by the appearance of the latter three absorption bands, which, in order from the highest wavenumber, we refer to as band I (orange filled area), band II (green filled area), and band (blue filled area), respectively. The center wavenumbers of the bands I—III were almost the same, but the relative intensity ratio was significantly different between human and bovine insulin fibrils. This observation suggests that the constituent iodine binding sites have very similar properties in both insulin fibrils but the number of binding site and/or the binding constants of polyiodides are different, which resulted in different colors.

### Structural propagation of insulin amyloid fibrils in self-seeding

We carried out selfseeding experiments to examine the propagation characteristics of amyloid structure. The spontaneously formed insulin fibrils analyzed in Figures 1 and 2 were defined as the 1st fibrils and used as seeds for the first run of self-seeding reaction. Afterwards, the seeding reactions were repeated using amyloid fibrils of the previous generation as seeds. As a result, ThT fluorescence intensity was preserved even after the repetitive self-seeding reactions in both species (Figure 3A), implying each fibril structure was replicated. The 1st fibrils showed lower intensity than the 2nd–5th fibrils, which suggests smaller number or lower affinity of ThT binding probably because the 1st fibrils tended to contain diverse fibril morphology (38) and fibril clumps with buried surfaces due to the formation at high temperature. The results of ATR-FTIR spectra and their second derivatives also showed the preservation of secondary structure even after the repetitive self-seeding reaction (Figure 3B, C).

**Figure 3.**
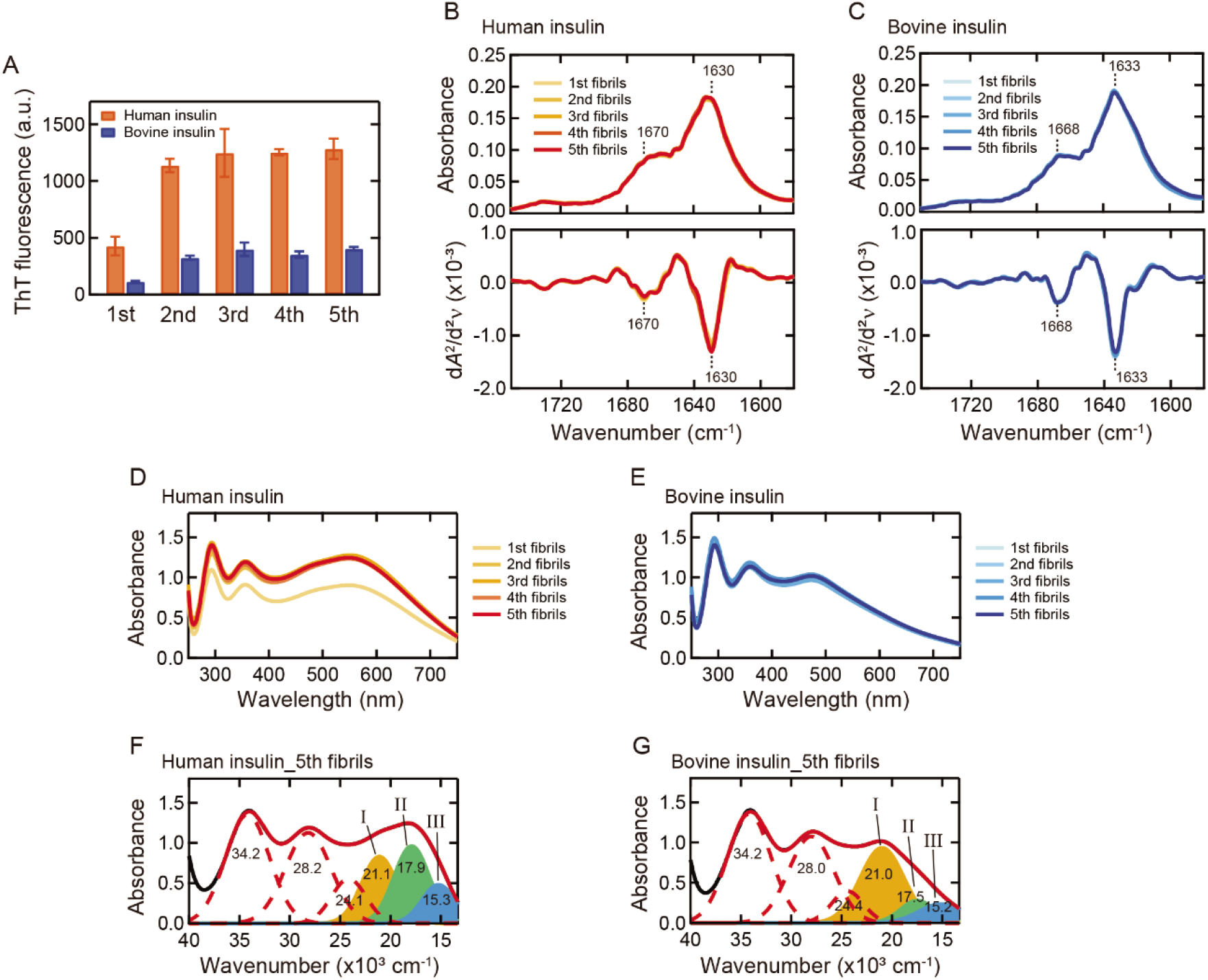
Self-seeding of human and bovine insulin. (A) ThT fluorescence intensity of the 1st-5th amyloid fibrils formed by repeats of self-seeding. The 1st fibrils correspond to the spontaneously formed amyloid fibrils analyzed in Figures 1 and 2 and were used as original seeds in this analysis. All samples were assayed in triplicate and error bars depict S.D. (B, C) ATR-FTIR absorption spectra of the 1st-5th amyloid fibrils (upper panel) and their second derivatives (lower panel) in amide I region of human (B) and bovine insulin (C). The spectra of the original seeds (i.e., 1st) are also shown. The positions of two main peaks of β-sheet and β-turn structures are indicated in the graphs. (D, E) UV-Vis absorption spectra of iodine-stained human (D) and bovine insulin fibrils (E) formed by self-seeding. For each absorption spectrum, the spectrum of unstained fibrils was subtracted to obtain net spectrum derived from iodine molecules. (F, G) Deconvolution of UV-Vis absorption spectra of iodine-stained final products (i.e., 5th) of human (F) and bovine insulin fibrils (G) obtained by self-seeding.

The replication of amyloid structure was further verified by iodine staining, which is expected to detect even slight structural differences with high sensitivity (29). The self-seeded 2nd-5th fibrils consistently exhibited the same shape of UV-Vis absorption spectra throughout all generations (Figure 3D, E), and as a consequence, the spectral deconvolution of the UV-Vis absorption spectra of the 5th fibrils reproduced almost the same absorption bands as those in the 1st fibrils (Figure 3F, G). The spectral intensity of the 1st fibrils was slightly lower than those of the 2nd-5th fibrils, especially in human insulin, although it was not so prominent as that in the case of ThT fluorescence intensity.

### Structural propagation of insulin amyloid fibrils in cross-seeding

We further carried out cross-seeding experiments to examine the propagation characteristics of amyloid structure between proteins with different amino acid sequences. In this work, human/bovine insulin proteins were cross-seeded with bovine/human insulin seeds, and the 2nd fibrils obtained in the self-seeding reaction (see Figure 3) were used as seeds for the first cycle of the cross-seeding. As a result, unlike self-seeding, ThT fluorescence intensity of the cross-seeded insulin fibrils gradually changed as generation progressed. Although the 1st fibrils showed similar intensity to that of the original seeds, ThT fluorescence intensity was gradually decreased in bovine insulin seeded with human insulin seeds, (Figure 4A, orange bars), meanwhile it was gradually increased in human insulin seeded with bovine insulin seeds (Figure 4A, blue bars). It was suggested that the templating ability of the original seeds is weak upon cross-seeding and thus allows fibril structure to change towards a more thermodynamically stable one.

**Figure 4.**
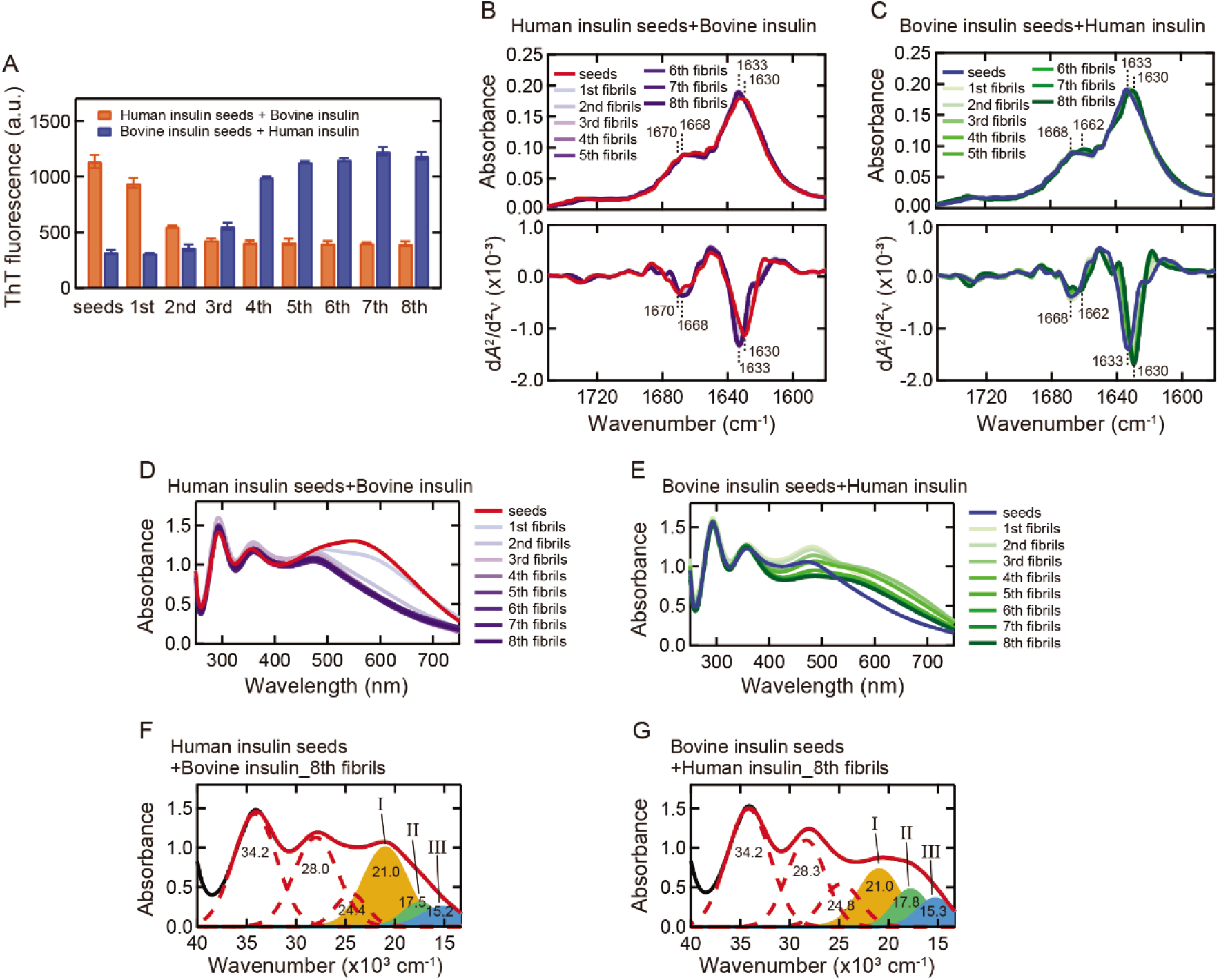
Cross-seeding of human and bovine insulin. (A) ThT fluorescence intensity of the 1st-8th amyloid fibrils. The intensities of the original seeds are also shown. All samples were assayed in triplicate and error bars depict S.D. (B, C) ATR-FTIR absorption spectra of the 1st-8th amyloid fibrils (upper panel) and their second derivatives (lower panel) in amide I region of bovine (B) and human insulin (C). The spectra of the original seeds are also shown. The positions of two main peaks of β-sheet and β-turn structures are indicated on the spectra of the seed fibrils and the 8th fibrils. (D, E) UV-Vis absorption spectra of iodine-stained bovine (D) and human insulin fibrils (E) formed by cross-seeding. (F, G) Deconvolution of UV-Vis absorption spectra of iodine-stained final products (i.e., 8th) of bovine (F) and human insulin (G) obtained by cross-seeding.

We next analyzed changes in secondary structure by ATR-FTIR spectroscopy. Similar to the ThT result, the spectra were gradually changed as the reaction was repeated (Figure 4B, C). When bovine insulin was seeded with human seeds, the spectral shape became similar to that of the bovine fibrils (Figure 4B, see also Figure 1E), suggesting that fibril structure changed to the intrinsic fibril structure of bovine insulin. Interestingly, when human insulin was seeded with bovine seeds, fibril structure did not change to the intrinsic human fibril structure. The spectral shape of the 8th fibrils did not coincide with that of human fibrils, and it showed a deeper second derivative minimum at 1630 cm^-1^ compared to that of human insulin fibrils and the position of the β-turn shoulder was shifted to 1662 cm^-1^ (Figure 4C, see also Figure 1E). It is estimated that a new-type of insulin fibrils with distinct β-sheet structure was formed. Far-UV CD analysis also showed a new spectral shape particularly when human insulin was seeded with bovine seeds (Figure S2).

We further tracked structural changes during the repeated cross-seeding by using iodine staining. The color of iodine staining detected structural changes successfully, and the UV-Vis absorption spectra of iodine-stained fibrils showed clear spectral changes in a longer wavelength region in both types of cross-seeding (Figure 4D, E). In agreement with the FTIR result, the absorption spectrum reached a distinct shape that was neither human nor bovine fibrils upon seeding of human insulin with bovine seeds, while it reached a shape similar to that of the bovine fibrils upon seeding of bovine insulin with human seeds. The spectral deconvolution of the 8th fibrils further verified differences in the composition of the bands IIII from those of the original seeds. When bovine insulin was seeded with human seeds, almost the same component of the absorption bands as those of bovine insulin fibrils was demonstrated (Figure 4F). When human insulin was seeded with bovine seeds, the relative intensity ratio was clearly different from those of human and bovine fibrils while the center wavenumbers were similar (Figure 4G).

### Morphology and cytotoxicity of the final fibril product after repeated self-seeding or cross-seeding

To further demonstrate the above-mentioned structural replication or change upon repeated self-seeding or cross-seeding, we performed morphological observation by TEM and AFM and cytotoxic assay. We selected the final generation of insulin fibrils formed by self-seeding (5th fibrils) and cross-seeding (8th fibrils) as samples. As a result, although it was difficult to identify any differences in appearance from TEM images (Figure 5A), AFM observation revealed the inheritance or change of fibril thickness successfully (Figure 5B). The thicknesses of self-seeded human and bovine insulin were 12.1 ± 2.0 nm and 6.9 ± 1.2 nm, respectively (*n* = 20), both of which were almost the same as those of the spontaneously formed human and bovine insulin fibrils. The thicknesses of the cross-seeded human and bovine insulin, on the other hand, were 15.1 ± 1.9 nm and 6.9 ± 1.3 nm, respectively (*n* = 20). The value of the cross-seeded bovine insulin supports the deviation to bovine-like fibril structure, and the thickest value of the cross-seeded human insulin strongly supports the formation of a new-type fibrils.

**Figure 5.**
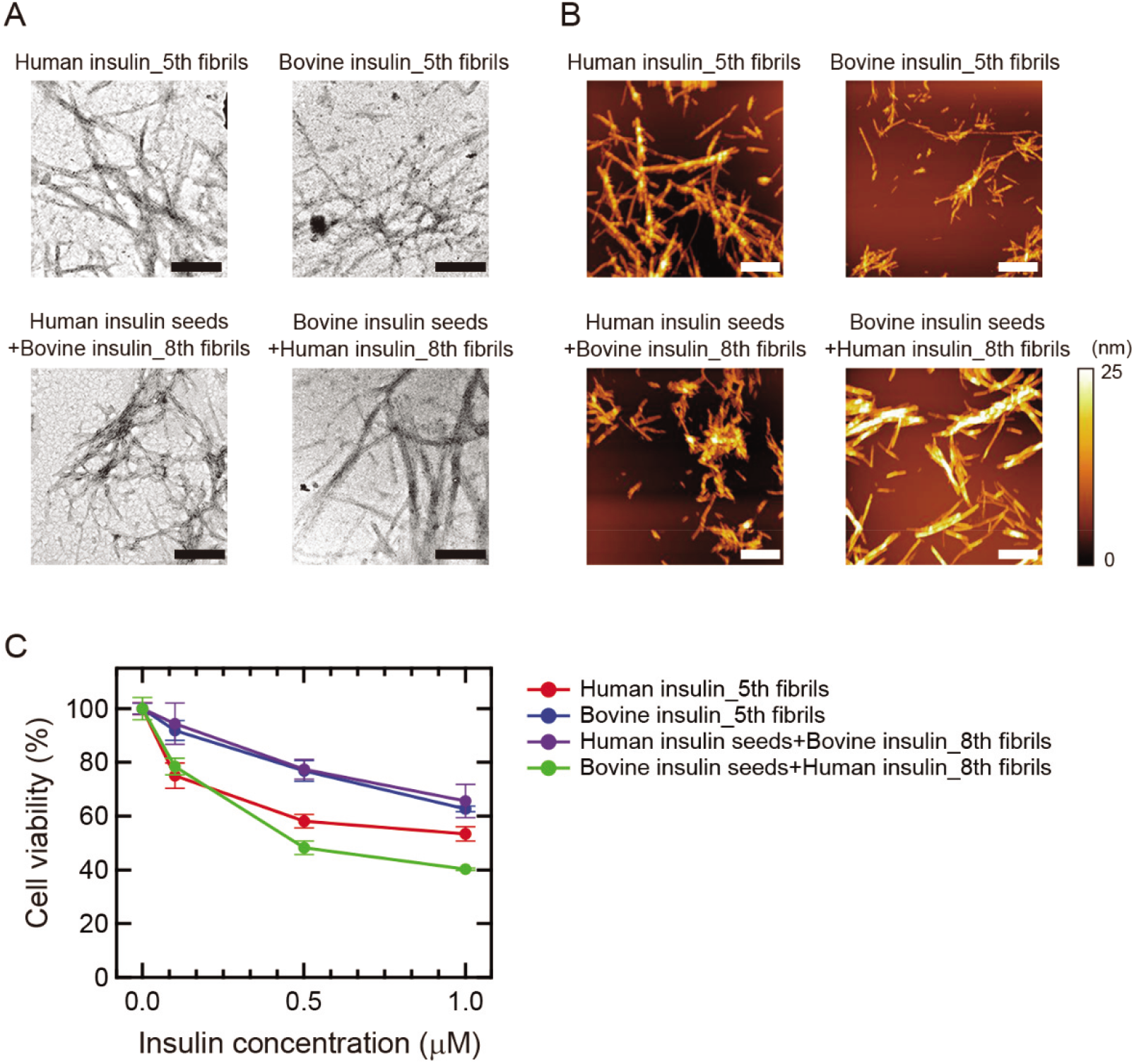
Morphology and cytotoxicity of final products of insulin fibrils formed by self- or cross-seeding. (A) TEM images. The black scale bars represent 200 nm. (B) AFM images. The white scale bars represent 1 μm. The color scale on the right represents the height of the sample. (C) Cell viability obtained from MTT assay of insulin fibrils at different concentrations using PC12 cells. Each sample was assayed in triplicate and the error bars depict S.D.

We next examined cytotoxicity against PC12 cells by MTT assay. As a result, all of the four types of insulin fibrils showed decrease in cell viability in a dose-dependent manner (Figure 5C). The cytotoxicity varied with the type of insulin fibrils, and the self-seeded human insulin yielded significantly lower values of cell viability than the self-seeded bovine insulin (Figure 5C, red and blue plots). This result indicates that the different structures of insulin fibrils have different cytotoxicity. The cross-seeded bovine insulin yielded almost the same cytotoxicity as the self-seeded bovine insulin (Figure 5C, purple plot), supporting the formation of bovine-like fibril structure. Interestingly, the cross-seeded human insulin yielded the lowest cell viability that was distinct from human or bovine fibrils (Figure 5C, green plot). This result indicates that new fibril structures emerged by cross-seeding have a potency to show a new phenotype.

### Investigation of the process of the structural changes during the repeated cross-seeding

To gain a detailed picture regarding the pathway of the structural changes induced by cross-seeding, SVD analysis was performed with the UV-Vis absorption spectra of iodine staining in the repeated cross-seeding shown in Figure 4D, E. A set of the UV-Vis spectra, ***M***, was decomposed as follows;

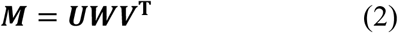

where ***U, W,V***^**T**^, represent the left-singular matrix, singular matrix, and right-singular matrix of the data-set matrix ***M***, respectively, assuming that ***M*** is a *m* × *n* matrix, ***U*** is a *m* × *m* unitary matrix, ***W*** is a *m* × *n* diagonal matrix, and ***V*^T^** is a *n* × *n* unitary matrix. In the analysis, we constructed ***M*** containing all the 18 spectra shown in Figure 4D, E in a manner where each column was composed of a single spectrum. In this case, each column of ***U*** is interpreted as a normalized diagonalized wave. The weights and proportions of the waves are described by the corresponding diagonal element of ***W***, which is called singular values, and corresponding row element of ***V*^T^**, respectively.

As a result, we obtained two dominant singular vectors, whose singular values satisfied accumulating contribution ratio of 0.94 (Figure 6A, B). With these singular vectors, which were referred to as SV1 and SV2 in contribution order, we carried out phase diagram analysis by plotting the ***V*^T^** values of SV1 versus those of SV2. Here, the result of the analysis using the whole wavelength range of the experimental data is shown. It was also confirmed that a similar result was obtained even when the wavelength range was limited to 400-750 nm, supporting that spectral changes in the visible region contribute importantly to this analysis. The phase diagrams obtained gave curved plots, suggesting multistep pathways in the both types of cross-seeding (Figure 6C, D). These plots were explained by assuming more than one liner segments, i.e., two liner segments in the cross-seeding of bovine insulin with human insulin seeds (Figure 6C), and four linear segments in the structural change in cross-seeding of human insulin with bovine insulin seeds (Figure 6D). Given that a single linear segment represents a two-state transition (39,40), the number of the lines corresponds to the number of transitions, and the line intersections in the diagrams correspond to metastable intermediate states routed in the process of cross-seeding.

**Figure 6.**
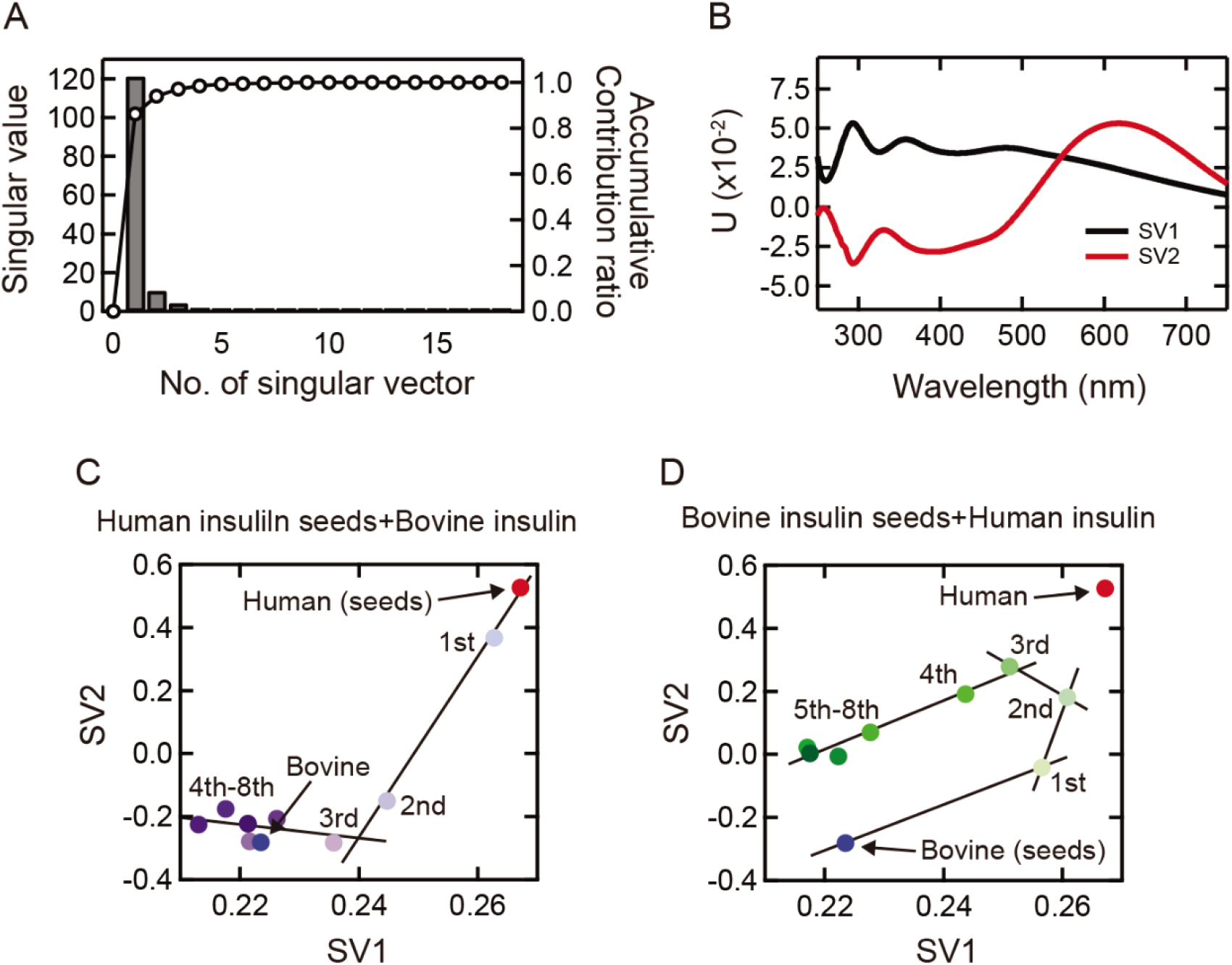
SVD analysis of UV-Vis absorption spectra obtained from iodine-stained insulin fibrils formed by cross-seeding. (A) The diagonal element of the singular matrix. (B) The 1st (SV1) and 2nd (SV2) columns of the left-singular matrix. (C, D) Phase diagrams obtained by plotting scores of the 1st (SV1) and 2nd (SV2) rows of the right-singular matrix. Solid lines, each of which represents a two-state transition, were superimposed to show that both of the plots can be described as a multistep transition.

## Discussion

### Structural changes induced by cross-seeding of human and bovine insulin

It was revealed that the cross-seeding of human and bovine insulin caused gradual structural changes in contrast to exact replication in self-seeding. The human and bovine insulin fibrils used in this study showed differences in structure in terms of fibril thickness, secondary structure and fibril surface (Figure 1, 2). Whilst repeated self-seeding preserved each of these structural properties strictly (Figure 3), significant changes from the original structure imprinted by seed fibrils were observed through successive generations of cross-seeding. This observation was similar to structural deviations reported previously for prion (25,41) and other proteins (26,27) (Figure 4), demonstrating weak ability to preserve original seed structures in cross-seeding.

The structural changes occurring in the repeated cross-seeding were successfully tracked by FTIR spectra and iodine staining, and in particular, the changes in visible color of iodine staining were obvious and clearly demonstrated complex changes. Iodine develops color by forming polyiodide ions in complex with crystalline compounds or polymers, and iodine-starch reaction is one of the best-known reactions in which iodine shows blue color by complexing with starch (42). Similarly, amyloid fibrils develop visible color with iodine, which is mechanistically explained by the formation of linear polyiodide ions composed of I_3_^-^ and I_5_^-^ on the fibril surfaces (29,43). Although the use of iodine staining in amyloid researches has not progressed so far due to the discovery of fluorescent dyes such as Congo Red and ThT, we have recently found that iodine staining could be used as a sensitive probe of structural polymorphism of amyloid fibrils (29). In this study too, iodine staining worked as a good strategy for evaluating structures of amyloid fibrils and their descendant generations formed by repeated self- or cross-seeding in detail. Considering that different color could be obtained even when the secondary structure was almost the same in our previous study (29), it is expected that the color of iodine staining senses higher-order structural differences like packing of protofilaments sensitively.

The most important finding obtained in this study is that the process of the repeated cross-seeding goes through one or more intermediate states. SVD analysis of the UV-Vis spectra obtained from the iodine-stained insulin fibrils revealed structural changes consisting of two or more structural transitions (Figure 6). Intriguingly, the number and types of intermediates varied depending on the direction of cross-seeding, and furthermore, a new fibril structure with distinct morphology and cytotoxicity appeared in a pathway-specific manner when human insulin was cross-seeded with bovine insulin seeds (Figure 5). The existence of metastable intermediate states is a strong experimental indication of the rugged energy landscape of amyloid fibrils, and the multistep structural changes as well as the pathway dependency of the convergent structure of amyloid fibrils are likely to be a natural consequence arising from the rough energy surface of the amyloid landscape that contains a number of local minima.

### Predicted scheme for propagation of amyloid fibrils in cross-seeding

On the basis of the present results, we propose a possible scheme of structural propagation induced by self-seeding and cross-seeding of human and bovine insulin. Figure 7A shows the case of bovine insulin. When bovine insulin is self-seeded, native bovine insulin reaches the inherent bovine-type fibril structure, which is presumed the most thermodynamically preferable fibril state of bovine insulin (Figure 7A, blue arrow). In contrast, upon cross-seeding, native bovine insulin first falls into the state of human-type fibril structure by templated with human insulin seeds (Figure 7A, red arrow). However, stepwise structural changes via an intermediate metastable state accompany and the fibril structure eventually adapts to the bovine-type fibril structure.

**Figure 7.**
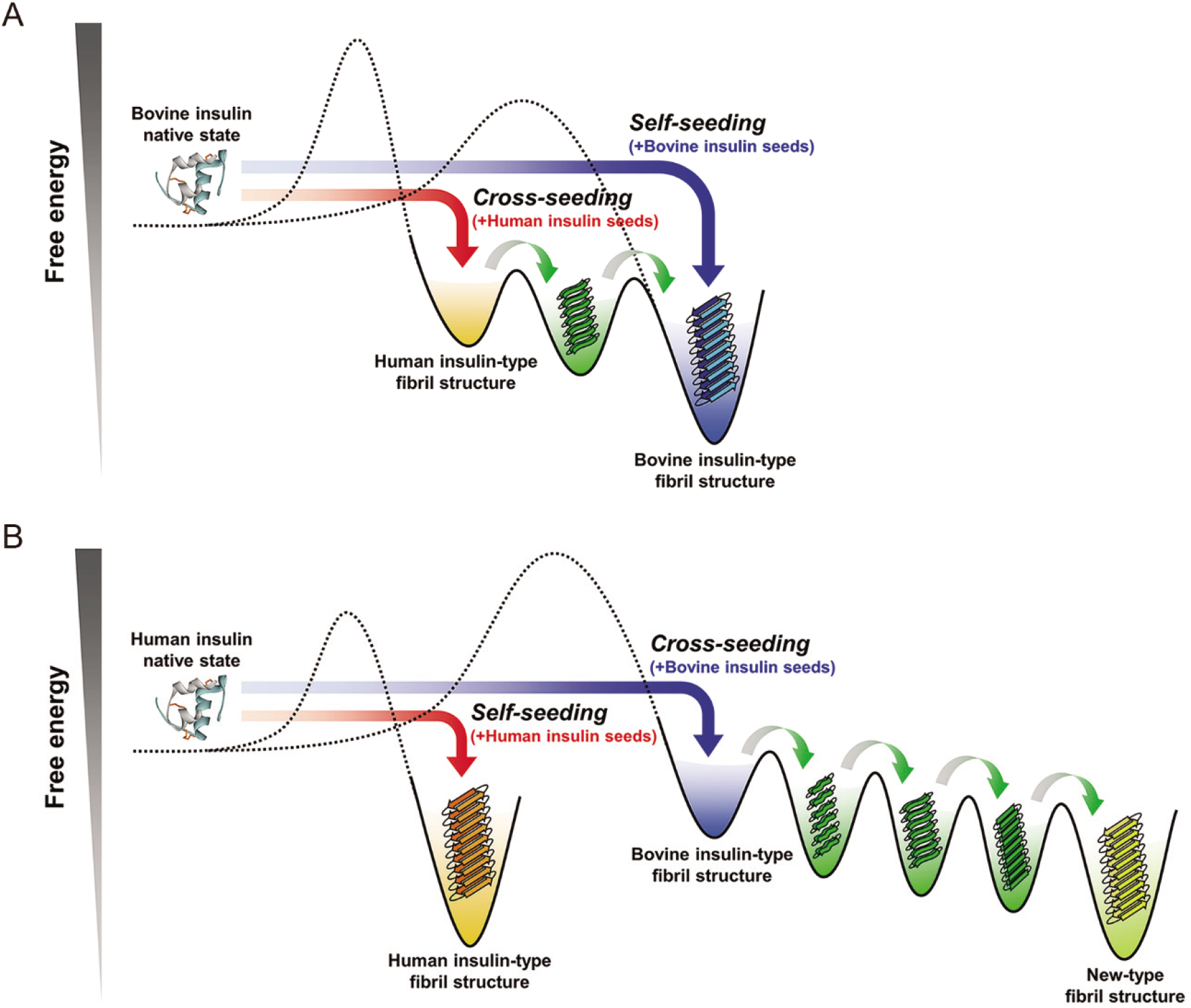
Schematic illustration for the energy landscapes of bovine (A) and human inulin (B). See the main text for details. It should be noted that the energy level of each structural state and the height of each energy barrier in the illustration do not include quantitative information.

On the other hand, in the case of human insulin (Figure 7B), the energy landscape is predicted to have different features, given that its cross-seeding eventually reaches a new-type fibril structure rather than the inherent human-type fibrils. When human insulin is self-seeded, native human insulin selectively reaches the inherent human-type fibril structure (Figure 7B, red arrow). When cross-seeded with bovine insulin seeds, native human insulin would first fall into the bovine-type fibril structure. However, the structure changes through several intermediate metastable states, and unlike the cross-seeding of bovine insulin in Figure 7A, the pathway leads to another fibril structure (Figure 7B, blue arrow). The intrinsic human insulin fibril structure is presumed to be blocked by a large energy barrier, resulting in the observation that human insulin finally took a new-type fibril structure.

Regarding the structural characteristics of the intermediate states, they could not be mentioned only from the results of iodine staining because the relationship between its color tone and the property of amyloid structure still remain unclear (29). To gain insights into the structural characteristics of the intermediates, we attempted SVD of ATR-FTIR spectra. Although the plots were more scattered than those of iodine staining, the phase diagram in the cross-seeding of human insulin with bovine seeds gave nonlinear shapes of the plots in agreement with the result of iodine staining (Figure S3). Additionally, the major peaks were observed in a range of 1620-1640 cm^-1^ in the singular vectors SV1 and SV2, from which it was implicated that the intermediate metastable states result from slight differences in β-sheet conformation.

### Implication for the manner of fibril propagation in cross-seeding

From previous studies of prion protein and other proteins, the “conformational selection and population shift” model, which has originally been proposed for explaining biomolecular recognition, has so far been recognized as a valid model to explain the molecular mechanism of cross-seeding (44). In this model, each protein is considered to possess a set of thermodynamically permissible structures, and the overlap of one or more of these structures between proteins with different sequences indicates the achievement of cross-seeding across a species barrier. Indeed, it can explain many experimental observations including cross-species prion transmission (41) and cross-talk between Aβ42 and Aβ40 (22). When this model is adapted to the energy landscape observed in this study, the thermodynamically permissible structures per protein correspond reasonably to a number of energy minima distributing over the landscape. If any of these energy minima coincide with the seed fibril structure, seed fibrils will interact with the counter protein molecules and guide them to fall to the energy minimum of the seed structure by skipping the nucleation barrier.

In addition to a large number of energy minima as candidates of fibril structures to be selected, an additional property worth noting is that some energy minima are positioned in adjacent on the landscape and tend to connect mutually by low energy barriers. This topology allows the deviation of fibril structure, so-called conformational switching, drift, adaptation, or evolution (25,45–47). In such cases, it is not so uncommon for fibrils to take a distinct structure from that of the original seeds as observed in the cross-seeding of the present study. Furthermore, it is expected that the energetic pathway to the final state will become rugged with multiple minima, resulting in the transient formation of metastable states. Considering this feature of the energy landscape, multiple cycles of cross-seeding will show dynamic equilibrium with transient accumulation of several different fibril species. If such energetic properties are generalized to a wide range of pathogenic proteins, cross-seeding will produce a variety of fibril structures over a time period of transmission, and thus, will be involved in pathology in a more complicated manner causing time-dependent fluctuation in infectivity and pathogenesis.

The energy landscape of amyloid fibrils is thought to vary with protein sequence (10), which will contribute to determine candidates of the final amyloid structure generated by cross-seeding. In addition to this, a specific interaction of protein molecules with seeds is another important factor, as it decides an initial landing point on the energy landscape (i.e., fibril templating). To investigate the role of the three different amino acid residues between human and bovine insulin in the latter issue, cross-seeding of human/bovine insulin was assessed using amyloid fibrils consisting of human A-chain or B-chain (48,49) as seeds. However, neither of these fibrils did not function as a fibril template, only accelerating fibril formation by shortening induction period (Figure S4). This result suggests that the interaction between the proteins and seeds is not restricted to local amino acid sequence, but more higher-order and complex factors are involved for exerting template ability.

### Conclusion

Through this study, a comprehensive understanding of the propagation mechanism of cross-seeding has been progressed from the perspective of energy landscape. The observation of the multistep structural changes has shed light on molecular details of the conformational deviations and the emergence of a new fibril structure. Remarkably, the presence of the metastable fibril states is a new physicochemical aspect of cross-seeding, and suggests the dynamic feature of amyloid structure during the time course of cross-seeding. The cross-seeding has been reported not only between isoforms of the same protein but also between different proteins, and it is considered to be important for understanding pathology and furthermore, cross-talk between different amyloid diseases (50–52). Given that heterologous proteins are actually involved in the pathology of several diseases, such as Alzheimer’s disease, prion disease, Parkinson’s disease and type 2 diabetes (22,24), accelerating researches on cross-seeding will pave new perspectives on the mechanism of pathology at the level of protein molecules.

## Data availability

All of the data described in the manuscript are contained within the manuscript.

## Acknowledgments

This work was performed in part under the Collaborative Research Program of Institute for Protein Research, Osaka University, CR −19-02. This work was supported by JSPS Core-to-Core Program, A. Advanced Research Networks.

## Funding and additional information

This work was funded by JSPS KAKENHI Grant Numbers JP16H04778, JP17H06352, and JP20H03224.

## Abbreviations

ThT: thioflavin T;
UV-Vis: ultraviolet-visible;
AFM: atomic force microscopy;
TEM: transmission electron microscopy;
ATR-FTIR: attenuated total reflection Fourier transform infrared;
SVD: singular value decomposition;
MTT: 3-(4,5-dimethylthiazol-2-yl)-2,5-diphenyltetrazolium bromide;
DMEM: Dulbecco’s Modified Eagle’s medium;
FWHM: full width at half-maximum.

